# LDL Receptor-Related Protein 5 is a Selective Transporter for Unesterified Polyunsaturated Fatty Acids

**DOI:** 10.1101/2023.01.16.524146

**Authors:** Wenwen Tang, Yi Luan, Qianying Yuan, Ao Li, Song Chen, Stanley Menacherry, Lawrence Young, Dianqing Wu

## Abstract

Polyunsaturated fatty acids (PUFAs), which cannot be synthesized by animals and must be supplied from the diet, have been strongly associated with human health. However, the mechanisms for their accretion and actions remain poorly understood. Here, we show that LDL receptor-related protein 5 (LRP5), but not its homolog LRP6, selectively transports unesterified PUFAs into a number of cell types. The LDLa ligand-binding repeats of LRP5 directly bind to PUFAs and are required for PUFA transport. LRP5 transports PUFAs via internalization to intracellular compartments including lysosomes, and n-3 PUFAs depends on this transport mechanism to inhibit mTORC1. This LRP5-mediated PUFA transport mechanism suppresses neutrophil extracellular trap formation in neutrophils and protects mice from myocardial injury during ischemia-reperfusion. Thus, this study reveals a previously unknown and biologically important mechanism for PUFA transport and mTORC1 regulation.

## Introduction

Fatty acids are classified into three major classes: saturated fatty acids (no double bonds), monounsaturated fatty acids (a single double bond), and polyunsaturated fatty acids (PUFAs) with multiple double bonds. Depending on the methyl terminal position of the first double bond, PUFAs are further divided into n-3 (or ω3) and n-6 (or ω6) PUFAs. The n-3 PUFAs include α-linolenic acid (ALA), eicosapentaenoic acid (EPA), and docosahexaenoic acid (DHA), whereas the n-6 PUFAs include linoleic acid (LA) and arachidonic acid (ARA). PUFAs are essential fatty acids, which cannot be synthesized by humans, mice and other mammals and thus must be supplied in the diet [1, 2]. PUFAs, particularly n-3 PUFAs, are generally considered to be anti-inflammatory and have beneficial effects on human health and particularly cardiovascular health including potential protection from myocardial ischemia-reperfusion injury [1–9]. While the molecular mechanisms accounting for these effects of PUFAs remain under-investigated, PUFA-derived bioactive metabolites regulate various aspects of inflammatory processes [10–12]. In addition, PUFAs can also signal through cell surface G protein-coupled receptors including GPR40 and GPR120 [13]. Moreover, studies showed that n-3 PUFAs inhibited mTORC1 signaling [14–18]. The mTORC1 signaling pathways regulate numerous important cellular processes including stimulation of protein synthesis primarily through phosphorylation of p70S6 kinase 1 (S6K), S6 protein, and eIF4E Binding Protein (4EBP)[19].

Although fatty acids can passively diffuse across the cell plasma membrane, cell surface transporters, including the CD36 class molecules and fatty acid transporter proteins (FATPs), are believed to facilitate the efficient uptake of fatty acids including PUFAs. However, they were either not rigorously tested for PUFA accretion or function, or only responsible for an esterified PUFA or under a special circumstance [20–25]. Thus, there is a general lack of knowledge on the mechanisms for PUFA accretion and the specific function of PUFAs in most tissues and cell types.

LRP5 is a single transmembrane protein and shares close amino acid sequence homology with LRP6. Its extracellular domain contains four YWTD repeat domains and three LDL receptor Class A (LDLa) or ligand-binding repeats [26]. The YWTD repeat domains are involved in interactions with Wnt proteins and Wnt antagonists including DKK and sclerostin [27], whereas no specific function has been linked to the LDLa repeats. These LDLa repeats are homologous with those in the LDL receptor that mediate lipoprotein binding [28]. LRP5 can function as a Wnt co-receptor in β-catenin stabilization via the binding of its intracellular domain to axin [29]. Studies, particularly mouse genetic studies, have further confirmed that LRP5 contributes to Wnt signaling, however, LRP6 seems to be the dominant co-receptor in Wnt-β-catenin signaling in most cases [30–34]. On the other hand, studies of LRP5 KO mice also reveal that some aspects of the LRP5 KO phenotype might not be readily connected to perturbation of Wnt-β-catenin signaling [35–40]. In this study, we identified LRP5 as being the first transporter selective for unesterified PUFA and showed that LRP5 is required for mTORC1 suppression by unesterified n-3 PUFAs, which is important for neutrophil function.

## RESULTS

### Neutrophil LRP5 protects mice from myocardial ischemia-reperfusion injury

Wnt-β-catenin signaling was previously implicated in neutrophil biology in the context of tumor immunity [41], and global LRP5-deficiency was shown to increase the extent of injury in a mouse model of myocardial infarction [42]. Thus, we investigated the roles of myeloid LRP5 and 6 in modulating injury during myocardial ischemia-reperfusion, a condition in which neutrophils play an important role [43, 44]. Myeloid-specific LRP5 (*Lrp5^M^*) or LRP6 (*Lrp6^M^*) knockout (KO) mice, driven by Lyz2-Cre, were subjected to left coronary artery ligation and reperfusion. The *Lrp5^M^* KO mice sustained significantly greater myocardial ischemia-reperfusion injury, as indicated by larger areas of myocardial infarction (normalized to the ischemic risk region produced by coronary artery ligation) and greater impairment in contractile function (as indicated by lower left ventricular ejection fractions) than those of the WT littermates (Fig. 1A & S1A). In contrast, myeloid-specific LRP6-deficiency mice had no significant differences in myocardial ischemia-reperfusion injury compared to their WT controls (Fig. 1B & S1A). LRP5-deficiency did not significantly alter the circulating neutrophil abundance prior to or after ischemia-reperfusion (Fig. S1B). However, the *Lrp5^M^* mice had moderately increased neutrophils, but not monocytes, in their hearts compared to WT, after ischemia-reperfusion (Fig. 1C & S1C,D). Moreover, there was significantly increased co-staining of MPO (a neutrophil marker) and citrullinated histone H3 (a NET marker) in hearts of *Lrp5^M^* KO mice vs. WT controls after ischemia-reperfusion (Fig. 1D & S1E), suggesting that myeloid-specific LRP5-deficiency increases neutrophil NET formation and probably activation. Consistent with this in vivo observation of increased formation of NETs in the LRP5-null hearts during ischemia-reperfusion, isolated LRP5-deficient neutrophils also had more citrullinated H3 histone than WT neutrophils upon stimulation (Fig. 1E) and produced more extracellular DNA than the WT cells (Fig. 1F).

**Figure 1.**
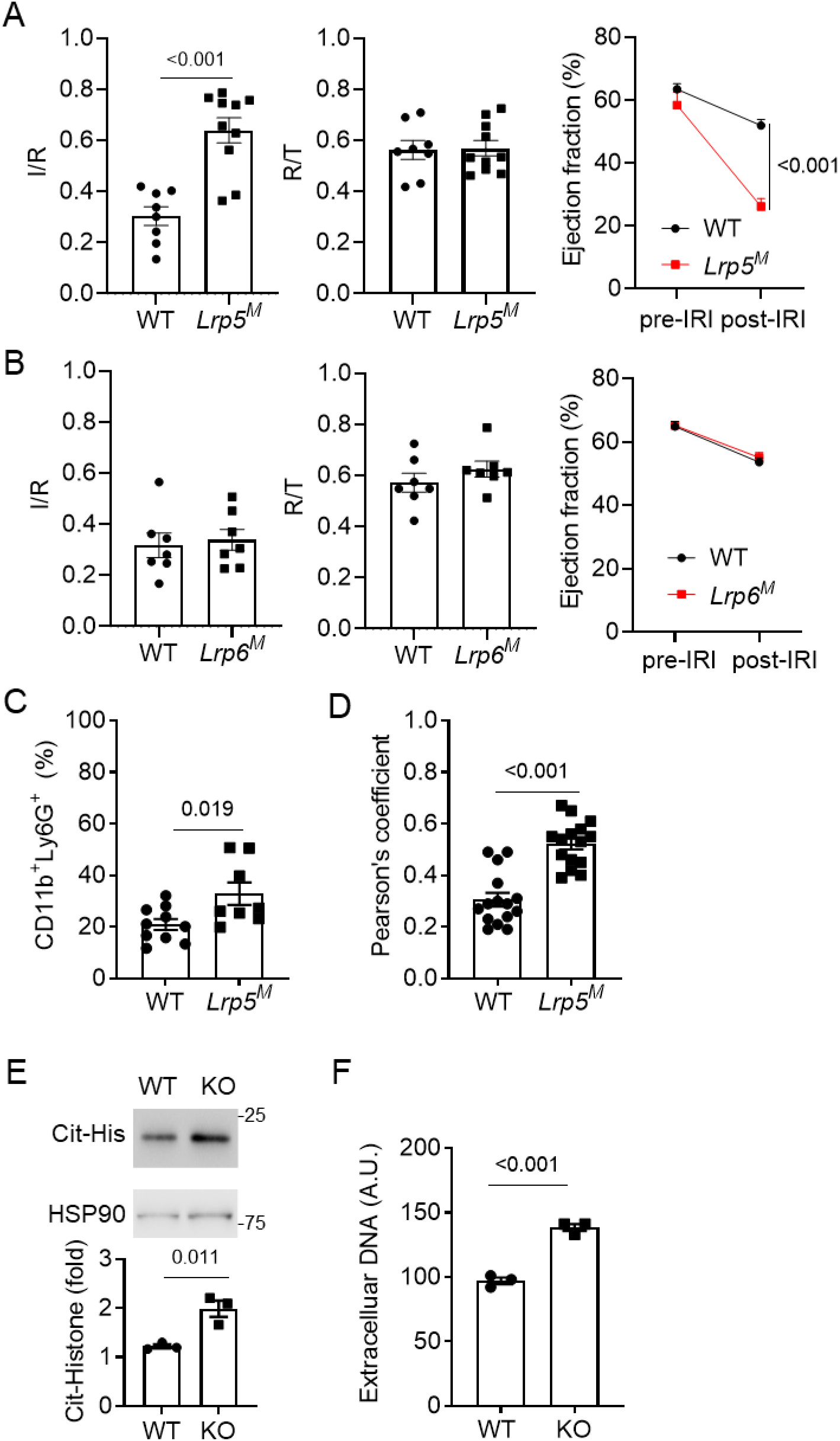
Myeloid-specific LRP5, but not LRP6, KO, leads to increased myocardial ischemia-reperfusion injury and NET formation. **A,B**) Mice lacking LRP5 (*Lrp5^M^*, **A**) or LRP6 (*Lrp6^M^*, **B**) in myeloid cells and their corresponding wildtype (WT) littermate control mice were subjected to myocardial ischemia-reperfusion injury. Infarction area (I) normalized against risk area (R) and Ejection Fractions before and after myocardial ischemia-reperfusion injury are shown. The ratios of risk area (R) to total area (T) are also shown. Each datum point represents one mouse**. C**) Neutrophil presence in injured hearts determined by flowcytometry. **D**) Pearson’s coefficients of co-localization of MPO and citrullinated histone (c-His) in the injured heart sections. Each datum point is one section. Three independent sections per mouse and five mice per group were analyzed. **E,F**) LRP5-deificient (KO) and wildtype (WT) neutrophils were, subjected to Western analysis to examine citrullinated histone H3 or ELISA to detect extracellular DNA. Data are presented as mean±sem with p values (Student’s t-test).

These results, together with the knowledge of the importance of neutrophils in myocardial ischemia-reperfusion injury, suggest that neutrophil LRP5 may be responsible for the observed phenotypes. Thus, we tested and found that Mrp8-Cre-driven LRP5 (*Lrp5^N^*) KO mice also had increased myocardial ischemia-reperfusion injury (Fig. 2A and S2A). Since Mrp8-Cre is more specific than Lyz2-Cre for neutrophils [45], these results indicate that neutrophil LRP5 specifically protects hearts from ischemia-reperfusion injury.

**Figure 2.**
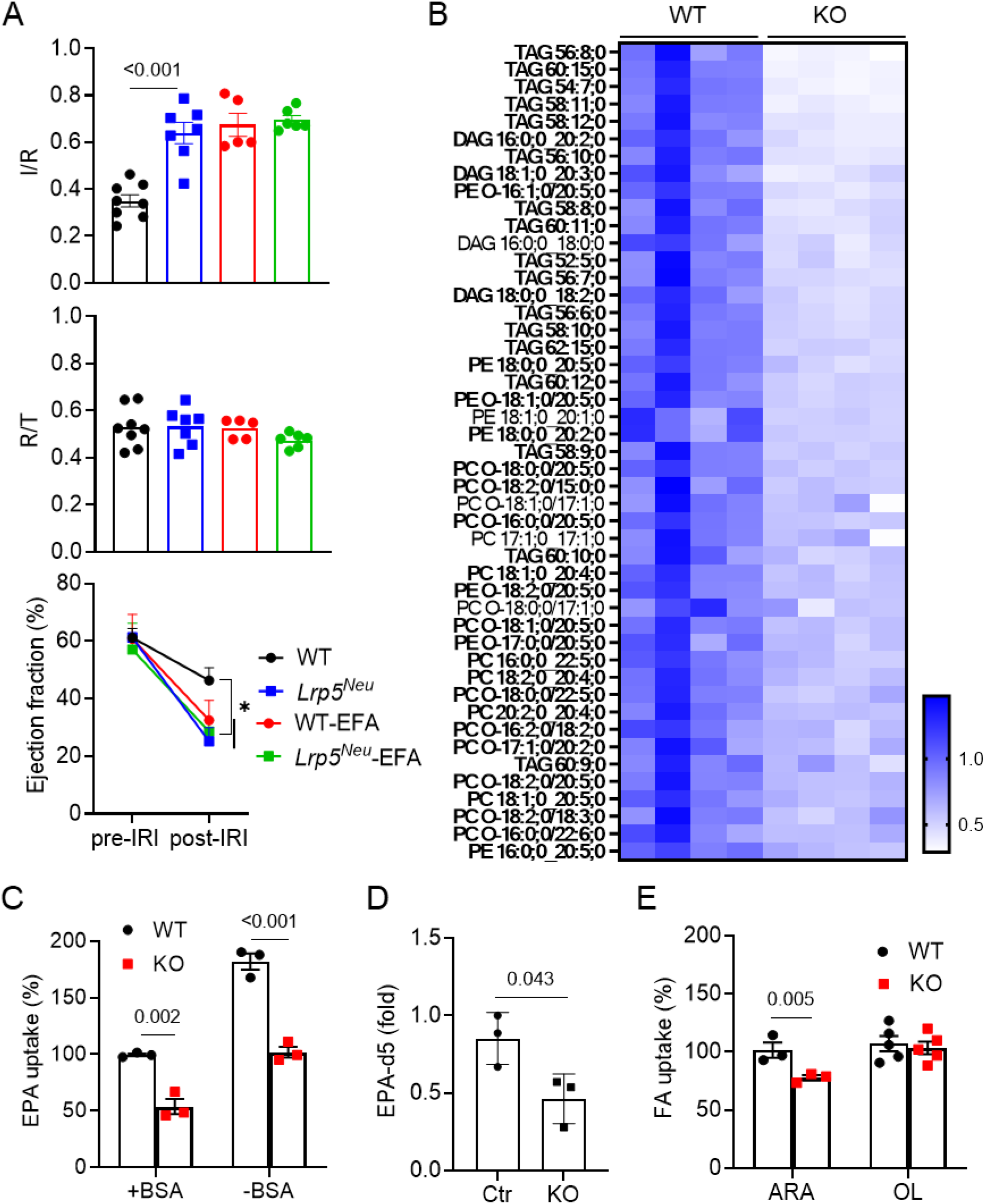
LRP5-deficiency reduces PUFA accretion in neutrophils. **A**) Mice lacking LRP5 in neutrophils (*Lrp5^Neu^*) and their corresponding wildtype (WT) littermate control mice on the normal diet (ND) or essential fatty acid-free diets (EFA) were subjected to myocardial ischemiareperfusion injury. Infarction area (I) normalized against risk area (R) and Ejection Fractions before and after myocardial ischemia-reperfusion injury are shown. Each datum point represents one mouse. Data are presented as mean±sem (Two-way Anova, * p<0.05). **B**) Heatmap of all of the glycerolipid species identified by the lipidomic analysis that show >0.7 Log_2_fold content reductions in LRP5 KO neutrophils compared to WT neutrophils with p<0.05. Those containing PUFAs are highlighted in bold. The complete dataset is shown in Supplementary Table 1. **C,E**) Neutrophils isolated from *Lrp5^Neu^* (KO) or WT mice fed on the essential fatty acid-free diet were incubated with ^14^C-EPA in the presence and absence of BSA or ^14^C-ARA or ^14^C-oleic acid (OL) with BSA. The uptake by WT cells is taken as 100%. **D**) Targeted LC-MS analysis of deuterated EPA in neutrophils. Data in C-D are presented as mean±sem (Student’s t-test).

### LRP5 is important for PUFA accretion in neutrophils

To understand the downstream signaling mechanism for LRP5 in neutrophils, we first examined the role of LRP5 in Wnt-induced β-catenin stabilization in neutrophils. We found that LRP5 deficiency had little effects on Wnt3a-stimulated β-catenin accumulation in neutrophils (Fig. S2B). In contrast, LRP6-deficiency abrogated Wnt3a-stimulated β-catenin accumulation in neutrophils (Fig. S2C). This result indicates that LRP6, rather than LRP5, mediates Wnt-β-catenin signaling in neutrophils. To identify the mechanism by which LRP5 functions in neutrophils, we performed lipidomic, transcriptomic, and metabolomic analyses of LRP5-null neutrophils in comparison to the WT cells. The lipidomic analysis revealed marked differences in the contents of glycerolipid species between LRP5-null and WT neutrophils (Supplementary Table 1). Predominantly, the lipids containing PUFAs were markedly reduced in LRP5-null neutrophils compared to WT (Fig. 2B). By contrast, LRP5 deficiency did not substantially alter the content of lipids that do not contain any PUFA (Fig. S2D). These lipidomic results were confirmed by targeted LC-MS analysis, which showed reduced contents of PC, PE and PA species containing PUFAs in LRP5-deficient neutrophils compared to WT neutrophils (Fig. S2E).

Because LRP5-deficiency significantly reduced PUFA accretion in neutrophils, we investigated the possibility that LRP5 might be involved in PUFA transport. Indeed, LRP5-deficiency resulted in a significant reduction in the uptake of ^14^C-EPA into neutrophils compared to the WT cells in the presence or absence of the fatty acid carrier protein bovine serum albumin (Fig. 2C). This result was further confirmed by detecting less deuterated EPA in neutrophils that were isolated from *Lrp5^N^* KO compared to those from control WT mice fed on the essential fatty acid-free diet and cultured with deuterated EPA (Fig. 2D). LRP5-deficiency also led to a reduction in ^14^C-ARA uptake by neutrophils (Fig. 2E). In contrast, LRP5-deficiency had no effect on ^14^C-oleic acid (OL, a mono-unsaturated fatty acid) uptake (Fig. 2E). These results together indicate that LRP5 is a PUFA transporter in neutrophils.

To determine if LRP5-transported PUFAs play an important role in neutrophil function *in vivo*, we examined the effects of PUFA-deprivation on myocardial ischemia-reperfusion injury. Because PUFAs cannot be synthesized and must be supplied from the diet, mice fed an essential fatty acid-free diet are deprived of PUFAs. In contrast to the previous findings in mice on a normal diet, the essential fatty acid-free diet abrogated the differences in myocardial ischemia-reperfusion injury between the neutrophil-specific LRP5 KO and control WT mice (Fig. 2A). Thus, the essential fatty acid-free diet replicated the effects of neutrophil-specific LRP5 KO on exacerbating myocardial ischemia-reperfusion injury in mice and had no additive effects in the LRP5 KO mice. These findings suggest that neutrophil PUFA depletion may account for the adverse phenotype of LRP5 KO mice during myocardial ischemia-reperfusion.

### LRP5 is important for PUFA accretion in other cell types

We went on to examine if LRP5 transports PUFA in other cell types. We performed the ^14^C-EPA and ^14^C-ARA uptake assay using NK cells isolated from Ncr1-Cre LRP5^fl/fl^, macrophages from Lyz-Cre LRP5^fl/fl^ mice, together with their corresponding LRP5 WT control cells. LRP5-deficiency resulted in significant reductions in ^14^C-EPA and ^14^C-ARA uptake in NK cells and macrophages (Fig. 3A). We also observed a decrease in ^14^C-EPA uptake by LRP5 KO HEK293 cells created by CRISPR/Cas9 in comparison with WT HEK293 cells (Fig. 3B). These results suggest that LRP5 transports PUFAs in cells other than neutrophils.

**Figure 3.**
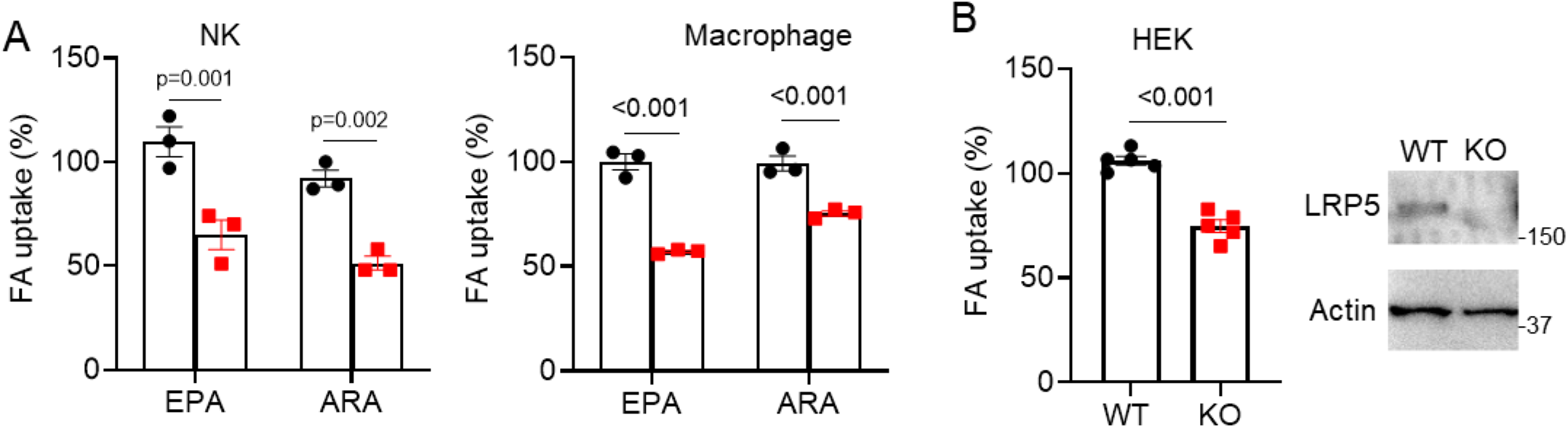
LRP5 deficiency reduces PUFAs uptake in various cell types. **A)** ^14^C-EPA and ^14^C-ARA uptakes were determined using splenic NK cells isolated from NCR-Cre LRP5 KO and WT littermates, peritoneal macrophages isolated from *Lrp5^M^* and littermate WT mice. **B)** ^14^C-EPA uptake was determined using LRP5 KO HEK293T cells and WT cells. LRP5 KO HEK293T cells were generated by CRISPR/Case9 and validated by Western analysis. Data are presented as mean±sem (Student t-test).

### LRP5 LDLa repeat domain binds to PUFAs

Because the LDLa repeats in the extracellular domain of LRP5 (Fig. S3A) are homologous with those in the LDL receptor that mediate lipoprotein binding and transport [28], we suspected that these repeats might be involved in PUFA transport. Indeed, when a LRP5 mutant lacking the LDLa repeats was expressed in the LRP5 KO HEK293 cells, there were no increases in ^14^C-EPA uptake (Fig. S3B). In contrast, overexpression of WT LRP5 increased ^14^C-EPA uptake (Fig. S3B). These data together suggest that the LDLa repeat domain is important for LRP5-mediated uptake of EPA. To determine if these repeats are involved in binding to PUFAs, we prepared conditioned medium (CM) from cells expressing a secreted form of LRP5 LDLa repeats fused with human IgG1 Fc region, designated as LDLa-Fc, and the control CM containing a secreted form of Fc (Fig. S3A). These CMs were added to a 96-well plate coated with BSA or BSA in the presence of one fatty acid. We found that CM containing LRP5 LDLa-Fc showed strong binding to EPA and DHA (Fig. 4A).

**Figure 4.**
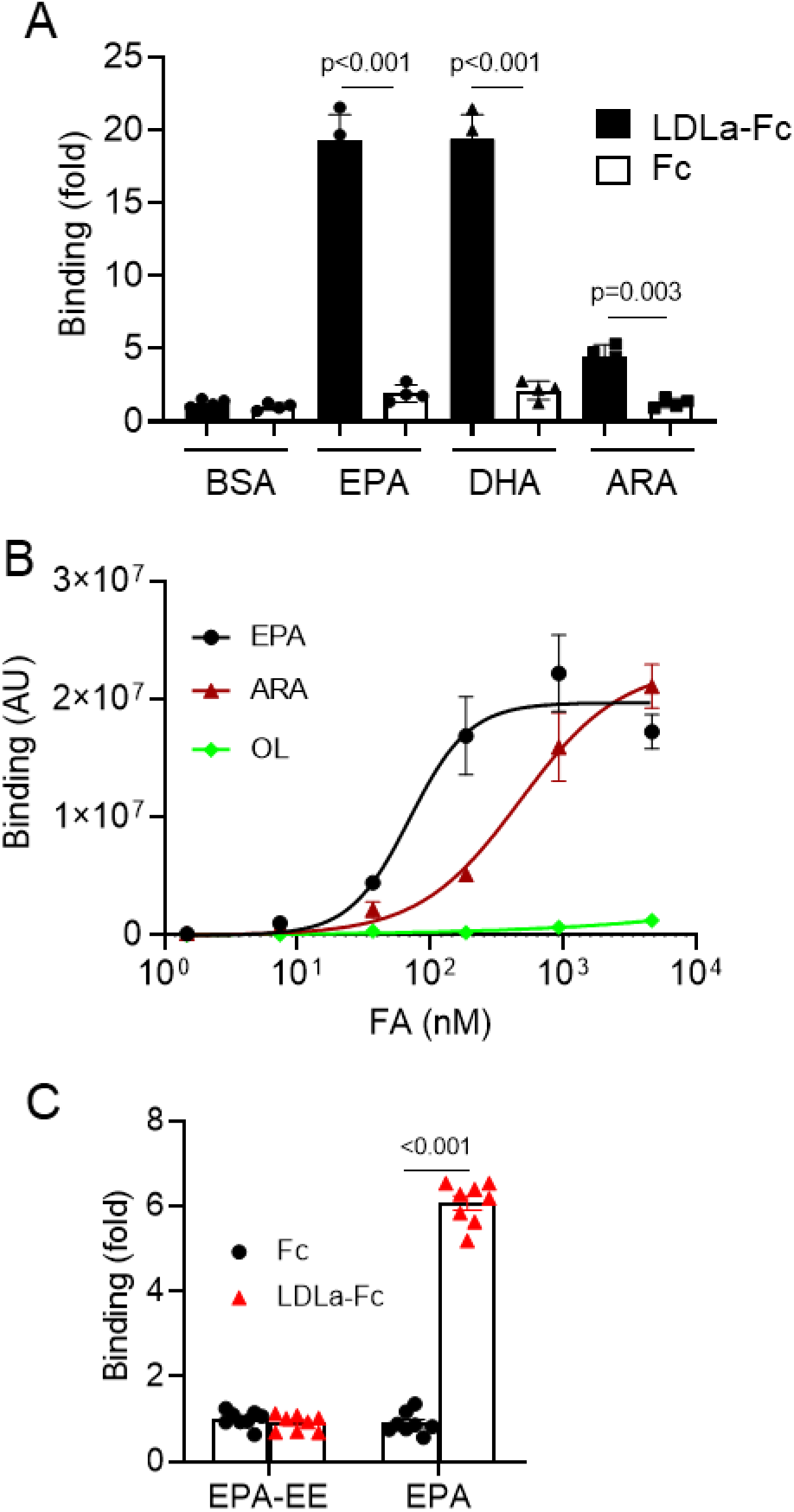
LRP5 LDLa repeats bind to PUFAs. **A**) Binding of LRP5 LDLa repeats to fatty acids was determined by coating the plate with BSA or BSA plus a fatty acid, followed with incubation with conditioned medium from cells expressing Fc or LRP5 LDLa-Fc. The binding of Fc to BSA is set as 1. **B**) Binding of purified LRP5 LDLa repeats fused with Fc to various fatty acids (non-linear fitting with Hill slope). Binding of these fatty acids to Fc was subtracted. AU, arbitrary unit. **C**) Binding of purified LRP5 LDLa repeats fused with Fc to EPA ethyl ester (EPA-EE). Data are presented as mean±sem with p values (Student’s t-Test).

To further characterize the binding affinities of LDLa-FC to the fatty acids, dose-dependent binding of purified LRP5 LDLa-FC to EPA, ARA and OL was performed (Fig. 4B). LDLa-Fc bound to EPA with a Kd value of ~69 nM and to ARA with a Kd value of ~479 nM (Fig. 4B), whereas binding to OL was barely detectable (Fig. 4B). These results are consistent with the binding results using CM. We also purified Fc-fused LRP6 LDLa repeats and showed no detectable binding to EPA (data not shown). Finally, we tested if LRP5 LDLa-Fc could bind to esterified PUFAs and found that it failed to bind to EPA ethyl ester (EPA-EE), an esterified form of EPA (Fig. 4C), suggesting that LRP5 may only transport non-esterified PUFAs.

### LRP5 transports EPA to intracellular compartments via internalization

Next, we examined the intracellular distribution of LRP5-transported EPA by staining neutrophils with an anti-PUFA antibody. This antibody preferentially recognizes EPA and DHA, but not palmitoylate, OL, or stearic acid (Fig. S4A). It also recognizes phosphatidylcholine containing EPA and DHA, but not myristic acid (Fig. S4B), suggesting that the antibody detects non-esterified PUFAs as well as PUFAs in phospholipids. When this antibody was used to stain neutrophils isolated from mice on the essential fatty acid-free diet, it produced a stronger signal in cells cultured with EPA than those without (Fig. 5A Panels *a* vs. *b*). PUFA antibody staining was partially colocalized with anti-TNG38 staining (a Golgi marker), suggesting that the antibody can be used to detect intracellular localization of loaded PUFAs in their unesterified and/or lipid forms. LRP5-deficiency markedly reduced anti-PUFA staining intensity, when normalized against TGN staining intensity, compared to that in WT cells (Fig. 5A, Panels *b* vs *c* & 5B), confirming that the anti-PUFA antibody detects LRP5-transported EPA. In addition to the localization at Golgi, LRP5-transported PUFAs were also detected at lysosomes, as LRP5-deficiency significantly reduced the colocalization of the anti-PUFA and anti-LAMP1 (a lysosomal marker) antibodies (Fig. 5C).

**Figure 5.**
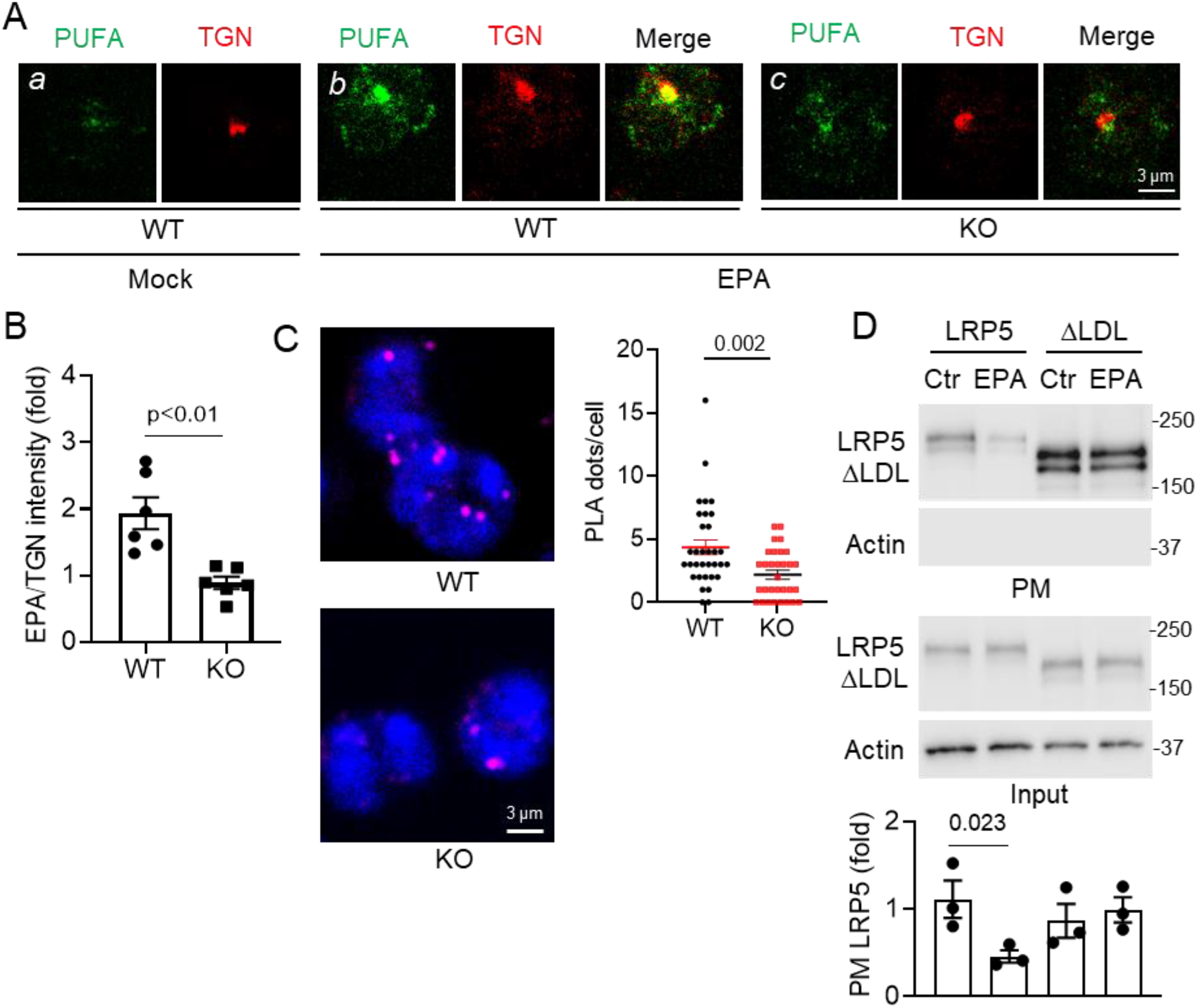
LRP5 transports PUFAs to intracellular compartments via internalization. **A,B**) Neutrophils from WT and neutrophil-specific LRP5 KO mice on the essential fatty acid-free diet and cultured with or without EPA (100 μM) for 3 hrs were stained with the anti-PUFA and anti-TGN38. Quantification of anti-PUFA staining intensity in Panels *b* and *c* with normalization against TGN staining intensity is shown in **B**. **C**) Neutrophils from WT and neutrophil-specific LRP5 KO mice on the essential fatty acid-free diet were cultured with EPA and subjected to proximity ligation assay (PLA) using antibodies for PUFA and LAMP1 (PLA fluorescence signal, magenta) and counterstained with DAPI (blue). **D**) LRP5 KO HEK293T cells were transfected with the LRP5 or LRP5 ΔLDL expression plasmid and treated with or without EPA. Cell surface LRP5 proteins were detected by Western blotting after precipitation of biotinylated cell surface proteins using avidin-columns. PM, plasma membrane. Data are presented as mean±sem with p values (Student’s t-Test).

The LDL receptor transport lipoproteins via internalization. We tested if EPA can promote LRP5 internalization using the LRP5 KO HEK293 cells re-expressing WT LRP5 or LRP5 lacking the LDLa repeats. We found that EPA reduced the amount of cell surface WT LRP5 without affecting the total LRP5 protein amounts (Fig. 5D). Consistent with the above observations, EPA did not reduce the amounts of surface LRP5 mutant lacking the LDLa repeats (Fig. 5D) or LRP6 (Fig. S4C). These results suggest that EPA promotes internalization of LRP5, but not LRP6, and that this action depends on the presence of LRP5 LDLa repeats.

### LRP5 is required for n-3 PUFA to suppress mTORC1

Next, we investigated mechanisms by which LRP5-transported PUFAs regulate neutrophils. n-3 PUFAs have been shown to regulate mTORC1 [14–18], and our IPA pathway enrichment analysis of differentially expressed genes revealed by total RNA sequencing of isolated bone marrow LRP5-null and WT neutrophils showed mTORC1 related signaling pathways (EIF2, eIF4-p70S6K, and mTOR signaling pathways) among the altered ones (Fig. S5A). Thus, we examined if LRP5 deficiency altered mTORC1 signaling activities. We found that phosphorylation of S6K and S6 was elevated in endogenously activated LRP5-null neutrophils isolated from inflammatory peritonea (Fig. 6A), as well as in naïve neutrophils isolated from bone marrow regardless chemoattractant stimulation (Fig. 6B), when compared to the corresponding WT control cells. There was a corresponding increase in the phosphorylation of 4EBP1, another mTORC1 downstream effector, in LRP5-deficient compared with WT neutrophils (Fig. 6A). By contrast, LRP6-deficiency did not affect S6K or S6 phosphorylation in neutrophils (Fig. S5B), and LRP5 KO did not affect phosphorylation of AKT, ERK, or p38 (Fig. 6B). In neutrophils, mTORC1 signaling regulates NET formation [46, 47]. Increased mTORC1 activity appeared to be responsible for increased NETs in the LRP5 KO neutrophils as LRP5-deficiency augmented histone citrullination (a NET surrogate marker) and this augmentation was abrogated by rapamycin (Fig. S5C).

**Figure 6.**
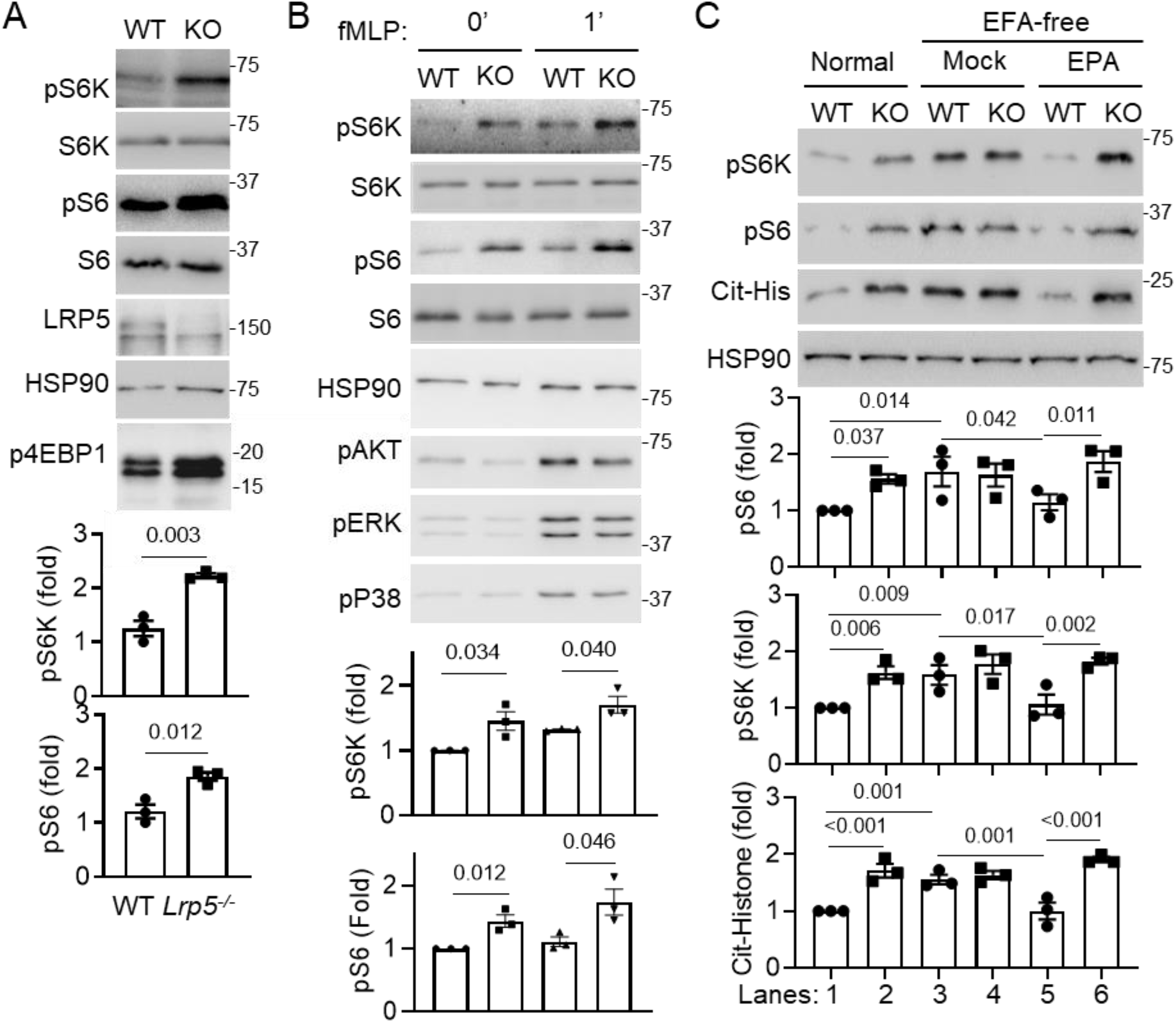
LRP5 is required for n-3 PUFA-mediated mTORC1 suppression in neutrophils. **A,B**) Western analysis of WT and LRP5 KO neutrophils isolated from peritonea pretreated with thioglycolate for 12 hrs (**A**) or bone marrow neutrophils stimulated with fMLP (1 μM, **B**). Western qualification was normalized to corresponding total proteins. **C**) Bone marrow neutrophils isolated from WT and neutrophil-specific LRP5 KO mice fed on the normal diet or essential fatty acid-free diet with or without gavage of EPA were analyzed by Western blotting. Western blot qualification was normalized to HSP90. Data are presented as mean±sem with p values (Student’s t-test for **A**,**B**; two-way Anova for **C**).

To determine if PUFAs underlie LRP5-deficiency-mediated activation of mTORC1 and NET formation in neutrophils, we examined the effect of PUFA-deprivation on mTORC1 signaling and histone citrullination. We found that PUFA-depleted WT neutrophils isolated from mice on the essential fatty acid-free diet had elevated S6K and S6 phosphorylation and histone citrullination compared to WT neutrophils isolated from mice on the normal diet (Fig. 6C, Lanes 1 vs 3). In contrast, PUFA-depletion had little effects on these events in LRP5 KO neutrophils (Fig. 6C, Lane 2 vs 4). Additionally, PUFA-depletion abrogated the differences between WT and LRP5-null neutrophils in these activities (Fig. 6C, Lanes 3 vs 4) that were observed between WT and LRP5-null neutrophils without PUFA-depletion (Fig. 6C, Lanes 1 vs 2). These results suggest that PUFAs may suppress mTORC1 signaling in an LRP5-dependent manner. To further corroborate this conclusion, mice on the essential fatty acid-free diet were given back a PUFA via gavage feeding of EPA, which decreased S6K and S6 phosphorylation and histone citrullination in WT neutrophils (Fig. 6C, Lanes 3 vs 5) without altering these events in LRP5 KO neutrophils (Fig. 6C, Lanes 4 vs 6). This LRP5-dependent inhibition of mTORC1 signaling by EPA was also observed in bone marrow neutrophils that were isolated from mice on the essential fatty acid-free diet and cultured with charcoal-filtered serum (deprived of fatty acids and lipids) (Fig. S5D). Specifically, EPA inhibited S6K and S6 phosphorylation and histone citrullination in the WT, but not LRP5-null, neutrophils. In addition, DHA also showed significant inhibition of S6K and S6 phosphorylation and histone citrullination in the WT, but not LRP5-null, neutrophils (Fig. S5D). ARA did not inhibit these mTORC1 signaling activities in neutrophils (Fig. S5E), which is consistent with previous observations that ARA does not inhibit mTORC1 signaling in other cell types [14–18]. These results together indicate that n3-PUFAs inhibition of mTORC1 signaling and NET formation in neutrophils requires LRP5.

We examined the effects of LRP5-deficiency on mTORC1 signaling in HEK293 cells. LRP5-deficiency also increased phosphorylation of S6K (Fig S6). Deprivation of PUFAs also increased S6K phosphorylation while abrogating the differences between WT and LRP5 KO in NK cells (Fig. S6). Importantly, EPA inhibited S6K phosphorylation only in WT cells, but not in LRP5 KO cells (Fig. S6), suggesting that EPA inhibition of mTORC1 in HEK293 cells also depends on LRP5..

## Discussion

In this study, we identified LRP5 as being a selective transporter for unesterified PUFAs that plays an important role in PUFA accretion in a number of different cells. Additionally, we showed that LRP5 was required for n-3 PUFA-mediated inhibition of mTORC1 in neutrophils and HEK293 cells. We also demonstrated the functional importance of LRP5-transported PUFAs in neutrophils; LRP5-transported PUFAs protect mice from myocardial ischemia-reperfusion injury. This function of LRP5-transported PUFAs is consistent with epidemiologic studies indicating that PUFAs, particularly n-3 PUFAs, are beneficial for heart health [1, 3–9].

While LRP5 had an important role for PUFA uptake in many different cells including neutrophils, NK cells and HEK293, it is not the only PUFA transporter in these cells. LRP5-deficiency showed partial inhibition of ^14^C-EPA and ^14^C-ARA uptake in these cells to varying degrees (Fig. 3). This is understandable, given the existence of other fatty acid transport mechanisms. The relative contributions of LRP5 to PUFA accretion in these cells varied probably due to the varying expression of different transporters. However, strikingly n-3 PUFA-mediated inhibition of mTORC1 signaling in these cells almost completely depended on LRP5 (Fig. 6C & S6), indicating that LRP5-transported PUFAs have distinct functions from those transported by other mechanisms. Because LRP5-mediated transport acts via LRP5 internalization (Fig. 5F), this mechanism can transport PUFAs to intracellular compartments including lysosomes in unesterified forms where unesterified PUFAs in turn inhibits mTORC1. By contrast, the mechanisms, such as CD36 and FATPs, transport fatty acids across plasma membranes. The fatty acids are subsequently bound to intracellular fatty acid-binding proteins (FABPs) and diffuse in the cytoplasm, and some may end up at intracellular compartments. Because addition of n-3 PUFAs failed to show significant inhibition of mTORC1 in LRP5 KO cells (Fig. 6C & S6), the non-LRP5 transported PUFAs may be either poorly targeted to lysosomes or largely esterified by the time they reach lysosomes. This explanation is further supported by the observed effect of LRP5 KO on PUFA presence in Golgi and lysosomes (Fig. 5A-C). Thus, this study for the first time demonstrates that different fatty acid transport mechanisms lead to different fates of transported fatty acids and distinct functional outcomes.

LRP5 is also distinguished from other known PUFA transporters for its selectivity to unesterified PUFAs. This molecular selectivity is likely due to the high affinity binding of LRP5 LDLa repeats for PUFAs compared with other fatty acids. LRP5 and LRP6 differ in the amino acid sequences of their respective repeats, which may also explain the inability of LRP6 to bind to and transport PUFAs. Of clinical interest in this regard, there is a potentially important human LRP5 polymorphism (rs3736228) at residue 1330 (Ala/Val) that is located in the second of the LDLa repeats. Although the association of this polymorphism with bone density/fracture is still up for debate, homozygous Val variants, which account for less than 5% of the population, seem to be associated with increased joint damage in patients with rheumatoid arthritis [48, 49]. Interestingly, the LDLa repeats containing this polymorphic Val residue showed reduced binding to EPA compared to the Ala-containing repeats (data not shown). Future studies are needed to determine if this polymorphism contributes to joint damage phenotypes via PUFA accretion.

Our study also provided examples for the functional importance of LRP5-transported PUFAs. Consistent with the epidemiological evidence of n-3 PUFAs being anti-inflammatory, LRP5-mediated accretion of n-3 PUFAs in neutrophils suppresses NET formation by inhibiting mTORC1 signaling. NETs are known to mediate tissue injury during inflammation including myocardial ischemia-reperfusion injury [50–56]. Thus, the increased mTORC1 signaling and NET formation likely exacerbated myocardial ischemia-reperfusion injury in the neutrophil-specific LRP5 KO mice. Although our results also indicate that COX2/lipoxygenase-mediated PUFA metabolites are unlikely to be involved in LRP5-mediated mTORC1 regulation (data not shown), they do not exclude the additional involvement of LRP5-transported PUFAs in other cellular effects. In addition, there could be mTORC1-independent actions of LRP5-transported PUFA in neutrophils, which would need to be further characterized in future studies. Knowing that LRP5-deficiency affects PUFA uptake in a broad range of cells, future studies also would need to investigate the importance of this LRP5 transport function in additional cell types for its contribution to LRP5 deficiency-related phenotypes.

## Methods

### Mice

The LoxP-floxed *Lrp5* (*Lrp5*^fl/fl^) and *Lrp6* (*Lrp6*^fl/fl^) mice were obtained from Bart Williams [57]. The *Lrp5*^fl/fl^ and *Lrp6*^fl/fl^ mice were back-crossed with C57BL/6N mice for more than seven generations before being intercrossed with Lyz2-Cre, Rosa26-CreER2, or Mrp8-Cre mice (Jackson lab), followed by additional backcrossing with C57BL/6N mice for more than ten generations. Ncr-1-Cre *Lrp5^fl/fl^* mice were described previously [35]. Wildtype C57BL/6N mice were purchased from Envigo. All animal experiments were performed with the approval of Institutional Animal Care and Use Committee (IACUC) at Yale University. Mice in all experiments were age- and gender-matched. Sample sizes used in animal experiments were based on the empirical determination in preliminary experiments. For deprivation of PUFAs, mice were placed on an essential fatty acid-free Diet (TD.84224, Envigo) for 4 days. For PUFA refeeding, 100 mg/kgbw PUFAs were delivered to mice 4 times via oral gavage every 12 hours just before experiments.

### Reagents and constructs

Antibodies to the following antigens were acquired commercially: Citrullinated-Histone-H3 (Abcam, ab5103), Myeloperoxidase/MPO (R&D, AF3667), LAMP-1 (Santa Cruz, sc-20011), TGN38 (Santa Cruz, Sc27680), PUFA (Cloud-Clone, PAO623Ge01), Beta-Catenin (BD, 610154), p-AKT473 (Cell Signaling Technology, 4060), mTOR (Cell Signaling, 2983), LAMP1 (Santa Cruz, sc-19992), Pacific blue-Ly6G (Biolegend, 127612), APC-CD11b (Biolegend, 101225), FITC-GZMB (Biolegend 515403), eFluor 450-IFNγ (ThermoFisher 12-7311-82), BV711-CD4 (Biolegend 100550), PE-CD8a (Biolegend 100708), BUV-395-CD45 (BD Biosciences 564279), PE-CD3 (Biolegend 100205), Alexa Fluor 488-CD27 (Biolegend 124221) Pacific blue-CD11b (Biolegend 101223), APC-NK1.1 (Biolegend 108710), Lrp5 (Cell Signaling Technology, 5731), Lrp6 (Cell Signaling Technology, 3395), Phospho-S6 (Cell Signaling Technology, 4858), Phospho-p70 S6K (Cell Signaling Technology, 9234), Phospho-4E-BP1 (Cell Signaling Technology, 2855), total 4E-BP1 (Cell Signaling Technology, 9452), Beta-Actin (Proteintech, 66009), and HSP90 (Proteintech, 13171). The following reagents were also acquired commercially: fatty acids were all from Cayman Chemical Company, rapamycin (Millipore Sigma, 553210), formyl-Met-Leu-Phe (fMLP) from Sigma (St. Louis, MO), Nordihydrogualaretic acid (Cayman Chemical, 70300), Celecoxib (Cayman Chemical, 10008672), GW9508 (Cayman Chemical, 10008907), and pFUSE-hIgG1-Fc2 (InvivoGen)

### Murine heart ischemia-reperfusion model

The mice were anesthetized with 5.0% isoflurane, intubated, and ventilated using a rodent ventilator. Anesthesia was maintained by inhalation of 1.5% to 2% isoflurane added to 100% oxygen. Body temperature was maintained at 37°C. The hearts were exposed. Ischemia was achieved by ligating the left anterior descending coronary artery (LAD) using an 8-0 silk suture with a section of PE-10 tubing placed over the LAD. After occlusion for 45 min, reperfusion was initiated by releasing the ligature and removing the PE-10 tubing. The chest wall was closed, the animal extubated, and body temperature was maintained at 37°C. Mice were re-anesthetized 24 hours later. The heart was surgically exposed, and the LAD was reoccluded by tying the suture in order to define the ischemic area at risk for myocardial infarction and the non-ischemic area. The non-ischemic myocardium was perfused with 1% Evans blue that was infused into the aorta in a retrograde fashion. Hearts were excised, sliced into five 1-mm cross-sections with the aid of an acrylic matrix (ZIVIC Labs). The heart sections were incubated with 1% triphenyl tetrazolium chloride solution (TTC, Sigma-Aldrich) at 37°C for 15 minutes. The non-ischemic area was stained blue, and within the ischemic region, residual viable myocardium was stained red and necrotic regions of myocardial infarction were unstained and appeared white. Using Image J software, the area of myocardial infarction was quantified and calculated as a percent of the myocardial area at risk, and the area at risk was calculated as a percent of the total LV area in each section.

### Neutrophil preparation

Murine neutrophils were isolated from bone marrows or peritoneal lavage. Briefly, bone marrow cells or peritoneal lavage cells (3% Thioglycolate induced overnight) collected from mice were treated with the ACK buffer (155 mM NH4Cl, 10 mM KHCO3 and 127 μM EDTA) for red blood cell lysis, followed by a discontinuous Percoll density gradient centrifugation. Neutrophils were collected from the band located between 81% and 62% of Percoll.

### Flow cytometry

Single-cell suspension was prepared, and the cell concentration was adjusted to 10^7^ cells/ml in staining buffer (1% BSA in PBS). Cells were pre-incubated with mouse Fc blocker (BD 553142, 1 μg/million cells in 100μl) on ice for 10 minutes, followed by incubation with fluorescent-labeled primary antibodies on ice for 30 minutes in the dark. Cells were then washed with staining buffer twice and resuspended in staining buffer for flow cytometry analysis.

### Internalization assay

HEK293T cells were treated with mock or fatty acids (100μM) in a culture medium for 3 hours. The cells were washed once with 1% fatty-acid-free BSA in PBS, followed by twice washing with ice-cold PBS. Cell surface proteins were biotinylated with 0.5mg/ml EZ-link-Sulfo-NHS-SS-Biotin (Thermo Fisher, A39258) in a PBS buffer with 2.5mM CaCl_2_ and 1mM MgCl_2_ on ice for 30 minutes. The reaction was stopped by the addition of PBS containing ice-cold 50 mM NH_4_Cl, followed by repeated washes with ice-cold PBS. The cells were then lysed in a buffer containing 1.25% Triton X-100, 0.25% SDS, 50 mM Tris-HCl, pH 8.0, 150 mM NaCl, 5 mM EDTA, 5 mg/ml iodoacetamide, 10 μg/ml PMSF, and the Roche proteinase inhibitor cocktail. After centrifugation, aliquots were taken as lysate controls, and the rest of the supernatants were used in pulldowns with NeutrAvidin beads (Thermo Fisher, 29200), followed by western blotting analysis.

### RNAseq and analysis

CD11b and Ly6G double-positive neutrophils were sorted out from the mouse bone marrow cells. Total RNA was isolated from the neutrophils using RNeasy Plus Mini Prep (Qiagen, 74134). The quality of RNA samples was measured using the Agilent Bioanalyzer. Then, RNA-seq libraries were prepared with the TruSeq stranded total RNA library prep kit (Illumina). single-end sequencing (75 bp) was performed on an Illumina HiSeq 2500 instrument at Yale Center for Genome Analysis. RNA-seq reads were aligned to the mouse reference genome (MM9) and gene expression was quantified by running the DESeq2 pipeline (default parameters). Differentially expressed genes with adjusted p-value less than 0.05 from each comparison were used for signaling pathway analysis in IPA (Ingenuity Pathway Analysis). The raw data are deposited at Gene Expression Omnibus (Accession number: pending).

### ^14^C-fatty acid Uptake assay

^14^C-labeled Eicosapentaenoic acid (0.2 μM, ^14^C-EPA, Moravek, MC2217), ^14^C-labeled arachidonic acid (2 μM, ^14^C-ARA, Moravek, MC364) or ^14^C-labeled oleic acid (1 μM, ^14^C-OL, American Radiolabeled Chemicals, ARC0297) was pre-mixed with or without 1% fatty-acid-free BSA (FAF-BSA) in RPMI1640 at 37°C for 1hr. Cells were washed once with serum-free medium, followed by incubation with FAF-BSA conjugated ^14^C-fatty acids at 37°C. After one hour, cells were first washed twice with 1% FAF-BSA in PBS and then washed with PBS once. Finally, cells were resuspended in PBS, followed with the scintillation cocktail and counted by a liquid scintillation counter (PerkinElmer, Tri-CARB 2100TR).

### Immunostaining of heart sections

Hearts were perfused and fixed with 4% PFA (Santa Cruz, sc-281692) for 4–6 hours on a shaker at 4 °C. They were then washed with PBS three times and perfused in 20% sucrose solution in PBS overnight at 4 °C. They were subsequently mounted in OCT embedding compound and frozen first at −20 °C and then at −80 °C. Tissue sections were prepared at 8-μm thickness with a cryostat and mounted onto gelatin-coated histological slides, which were stored at −80 °C. For immunostaining, slides were thawed to room temperature and fixed in pre-cold acetone for 10 min, then rehydrated in PBS for 10 min. The slides were incubated in a blocking buffer (1% horse serum and 0.02% Tween20 in PBS) for 1 - 2 hours at room temperature, then incubated with primary antibodies, which were diluted in the blocking buffer, overnight at 4 °C. The slides were subsequently washed three times with PBS and incubated with secondary antibodies diluted in blocking buffer for 1 hour at room temperature. After repeated washes, the slides were mounted with an anti-fade mounting media containing DAPI (Thermo Fisher, P36931) and visualized with a confocal microscope.

### Immunocytostaining

Primary neutrophils were then placed on poly-lysine-coated coverslips and fixed with 4% PFA for 10 min at room temperature followed by permeabilization with 0.03% saponin for 5 min at room temperature. After being rinsed with PBS three times, cells were blocked with a blocking buffer (2% BSA in PBS) for 1 hour at room temperature. Cells were then incubated with primary antibodies in blocking buffer at 4°C for overnight. The next day, secondary antibodies with conjugated fluorescent probes (Alexa488 colored in green and Alexa633 colored in red in the figures) were 1:200 in blocking buffer and incubated with cells for 1 hour at room temperature. Slides were prepared with the mounting medium containing DAPI and imaged under a confocal microscopy.

### Extracellular DNA measurement

Primary neutrophils were seeded into a poly-lysine coated 96-well plate at a density of 2×10^5^ cells/well and treated with 10 ng/ml GM-CSF for 4 hrs. 1000 mU/ml micrococcal nuclease (NEB, M0247S) was applied to cells in a volume of 100μl per well and incubated at 37°C for 10 minutes. The reaction was stopped with 5mM EDTA and the supernatant was collected and centrifuged at 200g for 8 minutes. The DNA in the supernatant was quantified using the Quant-iT Picogreen assay (Thermo Fisher, P7589).

### ELISA for anti-PUFA antibody and PUFA binding by LRP5 fragments

For detection of fatty acid binding by the anti-PUFA antibody, a 96-well white plate (NUNC polysorp, 437702) was coated with 1% fatty-acid-free BSA in PBS overnight at room temperature. The plate was rinsed once with PBS, followed by incubation with 100μl of 0.5 μg/μl fatty acid in PBS at 37°C for 1 hour. The plate was then washed three times with PBS and incubated with the anti-PUFA antibody (1:1500 dilution in TBST). After 2 hours later, the plate was rinsed with TBST for 3 times, followed by incubation with an HRP-conjugated goat anti-rabbit secondary antibody (Jackson ImmunoResearch, 111-035-144; 1:1000 dilution in TBST) for 30 minutes at room temperature. After three times washing with TBST, the plate was loaded with the SuperSignal pico plus chemiluminescent substrate (Thermo Fisher, 34580) and read by a plate reader (Perkin Elmer).

For detection of LRP5 fragment binding to PUFAs, a 96-well white plate was coated with 1% fatty-acid-free BSA in PBS overnight at room temperature. The plate was rinsed once with PBS, followed by incubation with a fatty acid at 37°C for 1 hour. The plate was then washed three times with PBS. The conditioned media or purified proteins were added to the plate. After 2 hours of incubation at room temperature, the plate was rinsed with TBST for 3 times, followed by incubation with an HRP-conjugated goat anti-human secondary antibody (Jackson ImmunoResearch, 109-035-003; 1:1000 dilution in TBST) for 30 minutes at room temperature. After three times washing with TBST, the plate was loaded with the SuperSignal pico plus chemiluminescent substrate (Thermo Fisher, 34580) and read with a plate reader (Perkin Elmer).

### Lipidomic analysis

For the analysis, quadruplicate samples were prepared from the *LRP5^N^* KO and WT control mice. Each sample consists of 10 million peritoneal neutrophils pooled from two mice. The cells were washed in PBS containing 1% fatty acid-free BSA for three times and snap-frozen in liquid nitrogen before being stored at −80 °C. The frozen samples were shipped on dry ice to Lipotype GmbH (Dresden, Germany), where they were extracted and analyzed using the Shotgun Lipidomic Platform.

### Targeted LC-MS/MS analysis of phospholipids

Cells (3-10 million) were collected and washed in PBS containing 1% fatty acid-free BSA for three times. Cell pellet was re-suspended in 600 μl of ice-cold chloroform/methanol (2:1, v/v) containing 1mM butylated hydroxytoluene (BHT) and incubated on ice for 30 minutes with occasional vortex mixing. Water (150 μl) was then added for 10 minutes. Samples were centrifuged at 2,000 rpm for 10 minutes at 4°C, and the organic layer (bottom layer) was collected to a new glass tube, whereas the aqueous phase (top layer) was re-extracted with 200 μl ice-cold chloroform/methanol (2:1, v/v). Organic phases from two-round extractions were combined and dried by a vacuum evaporator. Samples were reconstituted with 150ul methanol:H_2_O (2:1, v/v) containing 1% formic acid.

Phospholipids were separated using an Agilent 1200 HPLC system before introduction into a API 3000™ Triple Quadrupole Mass Spectrometer (Applied Biosystems). PC was separated by an Inertsil HILIC column and was acquired in the positive ESI mode, while PE and PA were separated by a C18 column and measured in the negative ESI mode. Collision energies ranged from 35 to 60 volts for various lipid species in MS/MS scan modes. Multiple reaction monitoring transitions of phospholipids were as follows: PC(O-36:5), 766.4>184.0; PC(36:5), 780.5>184.0; PC(38:5), 808.5>184.0; PC(O-38:5), 794.5>184.0; PA(18:0_20:5), 721.3>301.3; PA(20:1_20:5), 747.5>301.3; PE(P-18:0_20:5), 748.5>301.2; and PE(P-20:0_20:5), 776.5>301.2. Individual lipid species were quantified after normalization with spiked corresponding internal standards PC(14:0/14:0), PE(16:0-d9/16:0), and PA(16:0/18:1). **Targeted LC-MS/MS analysis of deuterated EPA.** Neutrophils were collected and washed in PBS containing 1% fatty acid-free BSA for three times. Cells were re-suspended in 100ul of PBS and then added with 100ul of 2.5N KOH/Meoh (1:4, v/v). Cell lysates were then incubated at 72°C for 15 minutes and acidified by adding 25ul formic acid followed by adding 225ul chloroform. The bottom chloroform layer was transferred to a glass vial and was dried. The dried extracts were reconstituted with 100ul chloroform:methanol (1:4, v/v) and separated by a C18 Column with mobile phases of 20mM Ammonium acetate (pH=9) and Methanol. Deuterium-labeled EPA was acquired in the negative ESI mode. Collision energies was 40 volts in MS/MS scan modes. Multiple reaction monitoring transitions for EPA-d5 was 306.2>262.2.

### Purification of LRP5 LDLa-Fc and Fc proteins

The Expi293 Expression System from ThermoFisher Scientific was used to express the Fc fusion proteins according to manufacturer’s instructions. The Expi293 cells were transfected using the ExpiFectamine 293 Transfection Kit. Protein expression in culture supernatant was confirmed 72 h post-transfection and harvested 7 days post-transfection. Cell debris and macrovesicles were removed from conditioned media by centrifugation at 380× g for 20 min at 4 °C. Recombinant human IgG1-tagged proteins were then purified by the Protein A/G agarose (Santa Cruz Biotechnology). Proteins were eluted with 100 mM glycine pH 3.0 and collected in tubes containing 1 M Tris buffer, pH 8.5. After dialysis with PBS overnight at 4°C, protein concentrations were determined by the Bradford assay.

### NK cell purification and culture

Mouse primary NK cells were isolated from the spleens of mice (8 weeks old) by using the NK cell isolation kits according to the manufacturer’s instructions (Miltenyi Biotec #130-090-864). Primary NK cells were cultured in RPMI-1640 (Gibco, 11875-093) supplemented with 10% FBS, penicillin (100 U/ml), streptomycin (100 μg/ml), 2-mercaptoethanol (500μM) and HEPES (10mM) at 37 °C supplemented with 5% CO2 in the presence of recombinant murine IL-15 (50 ng/ml).

### Proximity ligation assay

Primary neutrophils were placed on poly-lysine-coated coverslips and fixed with 4% PFA for 10 min at room temperature followed by permeabilization with 0.03% saponin for 5 min at room temperature. Cells were then incubated with pairs of antibodies (one from mouse and one from rabbit). PLA was carried out with the Duolink reagents from Sigma Aldrich according to the manufacturer’s directions. Briefly, cells were incubated with a pair of suitable PLA antibody probes in a humidified chamber, which were then subjected to ligation and amplification with fluorescent substrate at 37°C. The slides were mounted in mounting solution with DAPI. Cells were imaged with a Leica SP5 confocal microscope. Images were processed and quantified by Image J software as described [58].

### Statistical analysis and study design

Minimal group sizes for mouse studies were determined by using power calculations with the DSS Researcher’s Toolkit with an α of 0.05 and power of 0.8. Animals were grouped unblinded, but randomized, and investigators were blinded for most of the qualification experiments. No samples or animals were excluded from analyses. Assumptions concerning the data including normal distribution and similar variation between experimental groups were examined for appropriateness before statistical tests were conducted. Comparisons between two groups were performed by unpaired, two-tailed t-test. Comparisons between more than two groups were performed by two-tailed one-way ANOVA, whereas comparisons with two or more independent variable factors by two-tailed two-way ANOVA using Prism 9.0 software (GraphPad). Statistical analyses were based on biological replications. P<0.05 is considered as being statistically significant. All of the experiments were repeated at least twice, and representative ones are shown.

---

**Figure S1.**
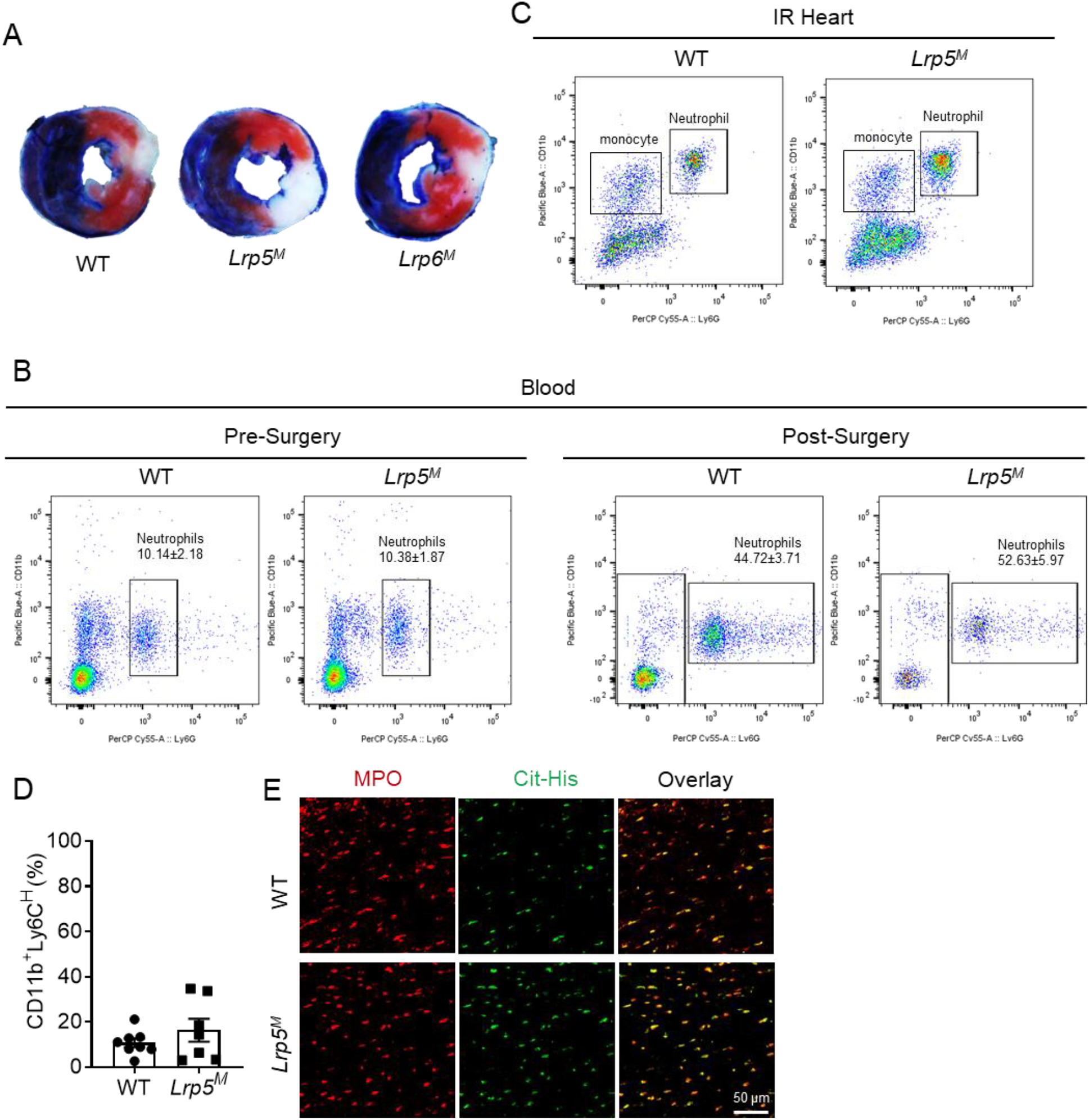
Myeloid-specific LRP5, but not LRP6, KO, leads to increased myocardial ischemia reperfusion injury and NET formation. **A**) Representative heart section images stained with triphenyl tetrazolium chloride from mice subjected to myocardial ischemia reperfusion injury for Fig. 1A,B. **B**) Flowcytometry analyses of blood neutrophils. Data are presented as mean±sem. No significance between WT and KO mice (two-tailed Student’s t-test, n>5). **C**) Representative flowcytometry charts for Fig. 1C. **D**) Monocyte presence in injured hearts determined by flowcytometry. **E**) Representative images of heart sections stained with anti-MPO and anti-citrullinated histone (Cit-His) for Fig. 1D.

**Figure S2.**
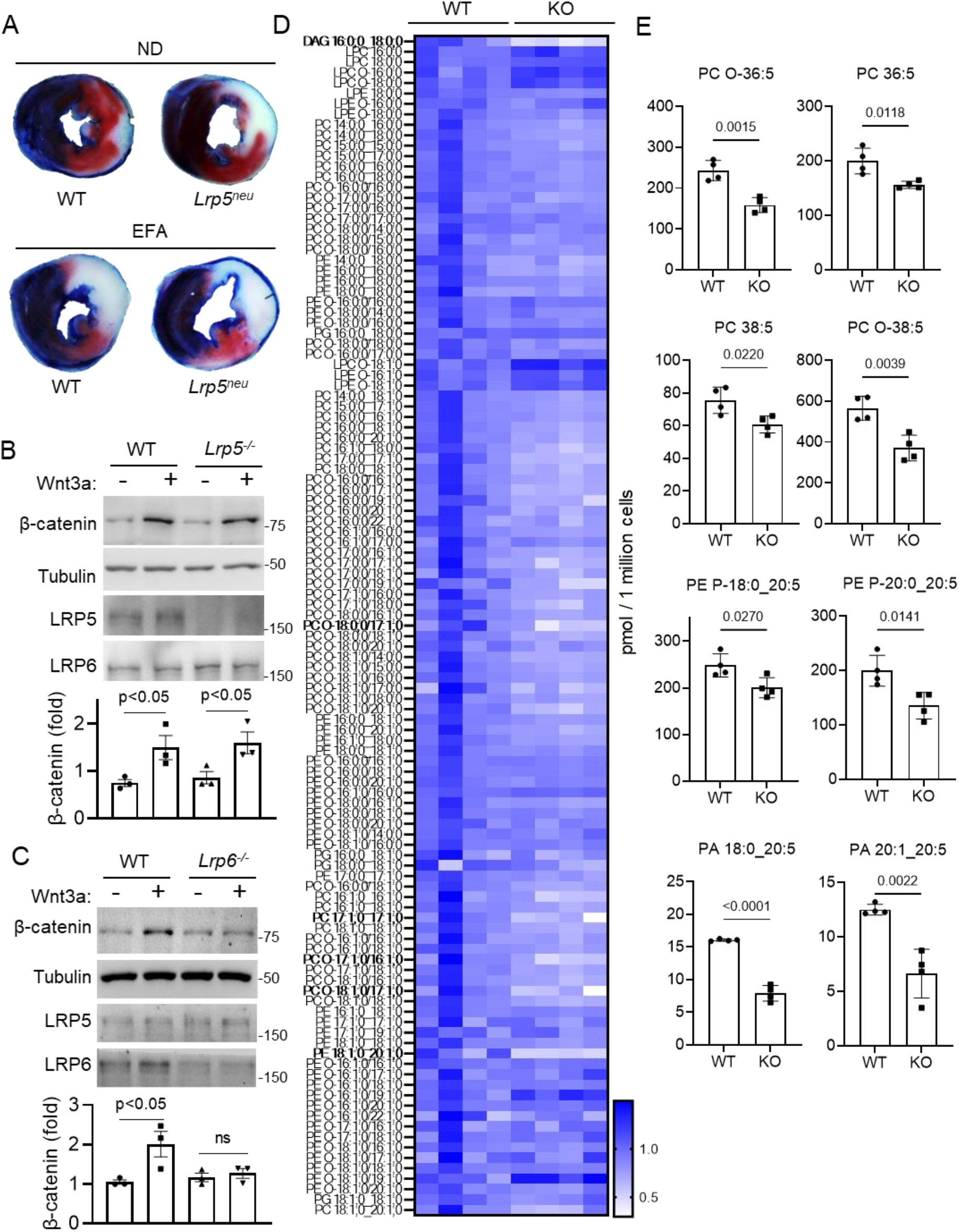
LRP5-deficiency does not affect Wnt-β-catenin signaling, but reduces PUFA accretion in neutrophils. **A**) Representative heart section images stained with triphenyl tetrazolium chloride from mice subjected to myocardial ischemia reperfusion injury for Fig. 2A. **B**) Bone marrow neutrophils were isolated from *Lrp6^M^* (**B**) or *Lrp5^M^* (**C**) and subjected to Wnt3A stimulation (50 ng/ml, 2 hours). **D**) Heatmap of all glycerolipid species identified by the lipidomic analysis that contain no PUFA. Those showing >0.7 Log_2_fold content reduction in LRP5 KO neutrophils are marked in bold. **E**) Targeted LC-MS analysis of PC, PE and PS species in neutrophils that were isolated from neutrophil-specific LRP5 KO or WT mice. Data are presented as mean±sem (Student’s t-test).

**Figure S3.**
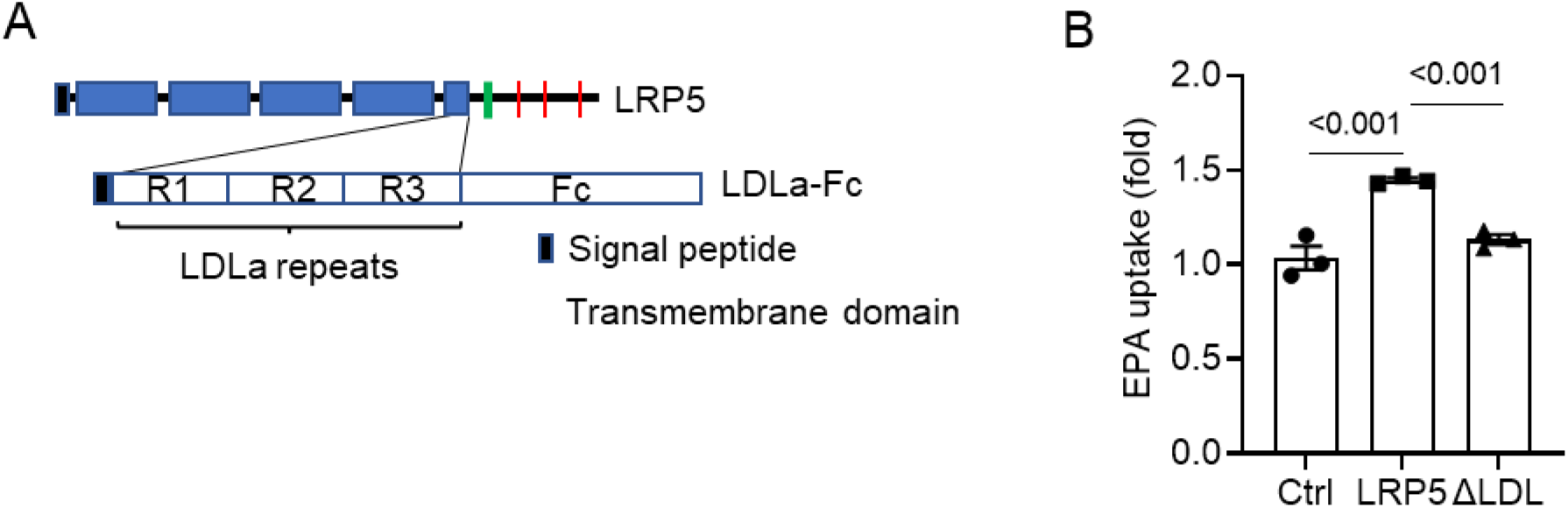
LRP5 LDLa repeats bind to PUFAs. **A**) Schematic representation of the full-length LRP5 protein and LRP5 LDLa-Fc fusion protein. **B**) LRP5 KO HEK293 cells were transfected with a control vector (Ctr) or a plasmid expressing mouse LRP5 or LRP5 without the LDLa repeats (ΔLDL). The cells were incubated with ^14^C-EPA with BSA. The uptake by the Ctr cells is taken as 1. Data are presented as mean±sem (One-way ANOVA).

**Figure S4.**
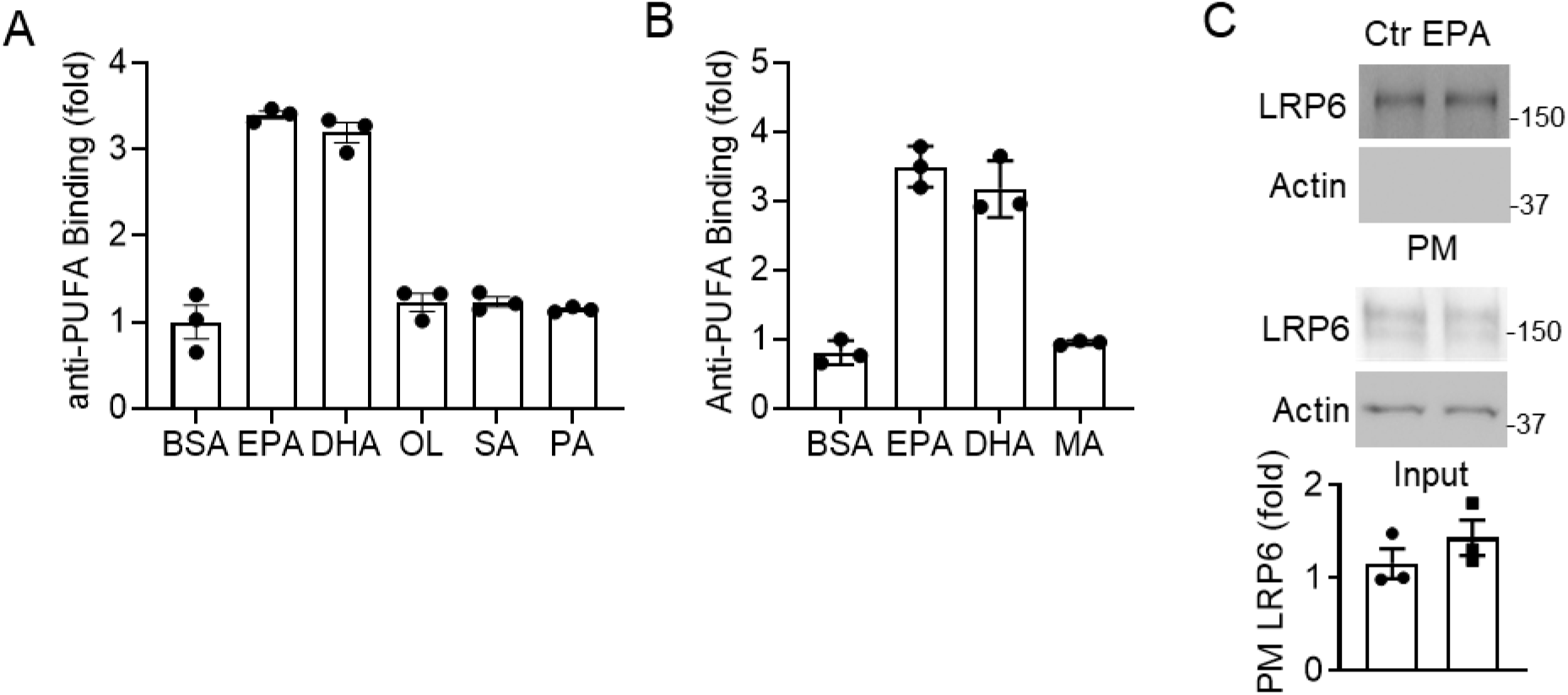
Anti-PUFA specificity and LRP6 internalization. **A**) Binding of anti-PUFA antibody to various fatty acids was determined by coating BSA or BSA carrying EPA, DHA, OL, SA (stearic acid) or PA (palmitoylate) in wells of a 96-well plate and incubating the anti-PUFA antibody. **B**) Binding of anti-PUFA antibody to phosphocholine (PC) containing EPA, DHA, or myristic acid (MA) was determined as in **A**. **C**) HEK293T cells were transfected with a LRP6 expression plasmid, and the internalization experiment was performed as in Fig. 4D. Data are presented as mean±sem.

**Figure S5.**
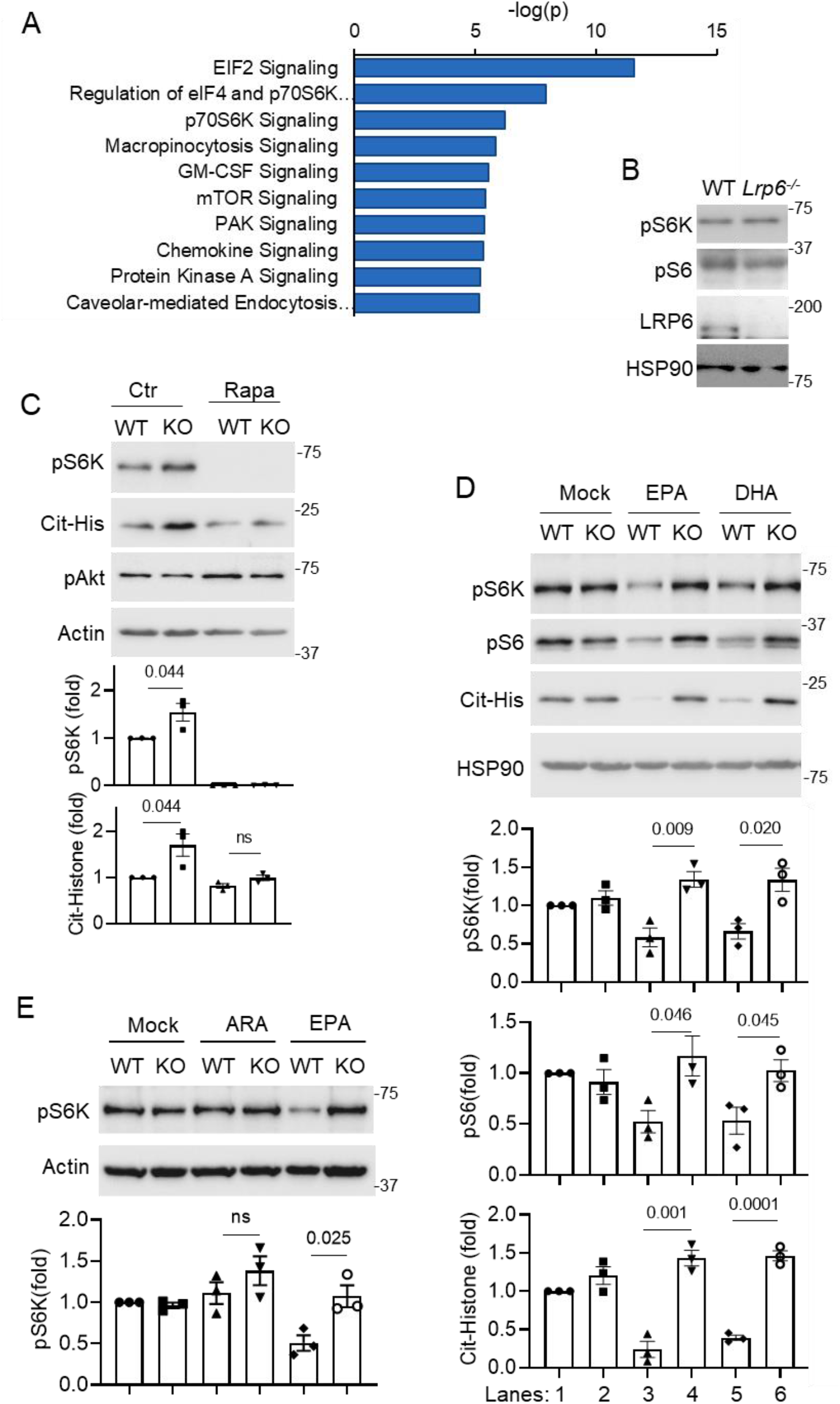
LRP5-transproted n-3 PUFA regulates mTORC1 signaling in neutrophils. **A**). IPA pathway analysis of differentially expressed genes based on transcriptomic analysis of isolated bone marrow LRP5-null neutrophils in comparison with WT neutrophils. **B**) Peritoneal neutrophils from WT and myeloid-specific LRP6 KO (*Lrp6^-/-^*) mice were analyzed by Western. **C**) Peritoneal neutrophils from WT and neutrophil-specific LRP5 KO mice were cultured with10 ng/ml GM-CSF for 4 hrs in the presence and absence of rapamycin (200 nM). Western qualification was normalized to actin. **D,E**) Neutrophils isolated from neutrophil-specific LRP5 KO or WT mice fed on essential fatty acid-free diet were cultured with mock or 100 μM n-3 PUFAs as well as 10 ng/ml GM-CSF in the charcoal-filtered serum medium for 3 hours before being analyzed by Western blotting. Western qualification was normalized to HSP90 or actin.

**Figure S6.**
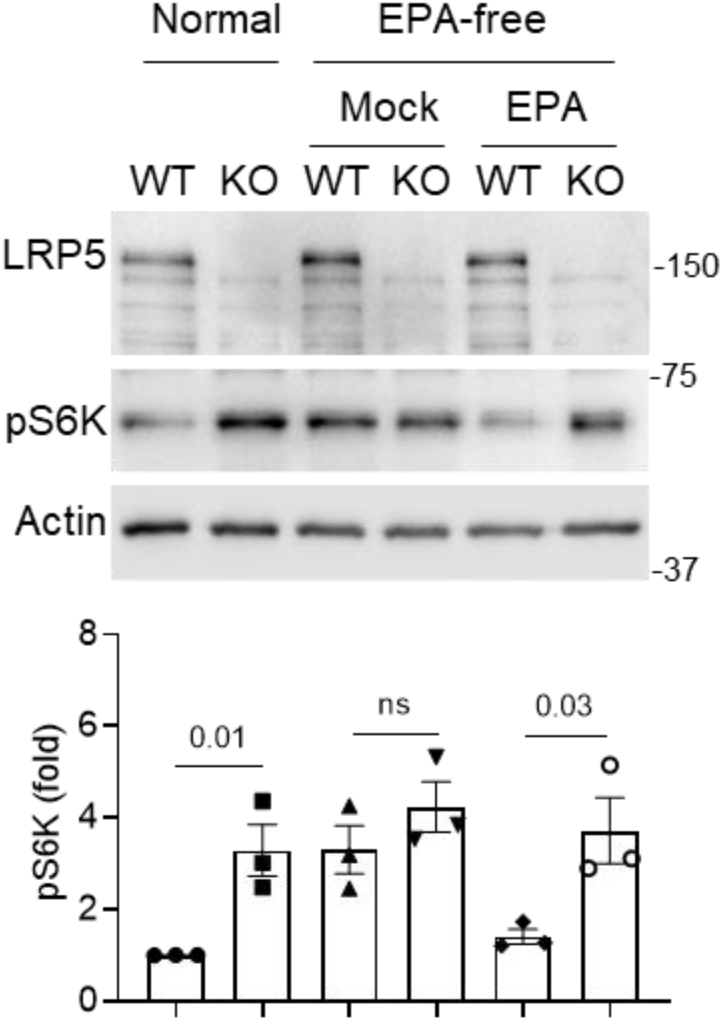
LRP5-transproted n-3 PUFA regulates mTORC1 signaling in HEK293 cells. WT and LRP5 KO HEK293 cells are cultured in the normal or charcoal-filtered serum for overnight, followed by being cultured in the presence or absence of EPA for 2hrs before Western blotting analysis. Data are presented as mean±sem with p values (Student’s t-test).

